# Immune-Based Prediction of COVID-19 Severity and Chronicity Decoded Using Machine Learning

**DOI:** 10.1101/2020.12.16.423122

**Authors:** Bruce K Patterson, Jose Guevara-Coto, Ram Yogendra, Edgar Francisco, Emily Long, Amruta Pise, Hallison Rodrigues, Purvi Parikh, Javier Mora, Rodrigo A Mora-Rodríguez

## Abstract

Individuals with systemic symptoms long after COVID-19 has cleared represent approximately ~10% of all COVID-19 infected individuals. Here we present a bioinformatics approach to predict and model the phases of COVID so that effective treatment strategies can be devised and monitored. We investigated 144 individuals including normal individuals and patients spanning the COVID-19 disease continuum. We collected plasma and isolated PBMCs from 29 normal individuals, 26 individuals with mild-moderate COVID-19, 25 individuals with severe COVID-19, and 64 individuals with Chronic COVID-19 symptoms. Immune subset profiling and a 14-plex cytokine panel were run on all patients. Data was analyzed using machine learning methods to predict and distinguish the groups from each other.Using a multi-class deep neural network classifier to better fit our prediction model, we recapitulated a 100% precision, 100% recall and F1 score of 1 on the test set. Moreover, a first score specific for the chronic COVID-19 patients was defined as **S1 = (IFN-γ + IL-2)/ CCL4-MIP-1β**. Second, a score specific for the severe COVID-19 patients was defined as **S2 = (10*IL-10 + IL-6) - (IL-2 + IL-8)**. Severe cases are characterized by excessive inflammation and dysregulated T cell activation, recruitment, and counteracting activities. While chronic patients are characterized by a profile able to induce the activation of effector T cells with pro-inflammatory properties and the capacity of generating an effective immune response to eliminate the virus but without the proper recruitment signals to attract activated T cells.

**Summary:** Immunologic Modeling of Severity and Chronicity of COVID-19

## INTRODUCTION

Chronic COVID-19 is a group of previously infected individuals, so called “Long Haulers”, who experience a multitude of symptoms from several weeks to months after recovering from their acute illness and presumably months after viral clearance. These symptoms include joint pain, muscle aches, fatigue, “brain fog” and others. These symptoms can commonly resemble rheumatic diseases such as rheumatoid arthritis, autoimmune disorders, and others such as fibromyalgia and chronic fatigue syndrome (1). Many of these common disorders are caused by inflammation, hyper- and/or auto-immunity and some such as chronic fatigue are associated with viral persistence after an acute infection with pathogens such as Epstein Barr and Cytomegalovirus (2). Recent studies including those from our laboratory have suggested that (CC) may be caused by persistent COVID itself (3). Here, we sought to identify possible immunologic signatures of COVID-19 severity and to determine whether Chronic COVID-19 might represent a distinct immunologic entity compared to mild to moderate (MM) or severe/critical COVID-19. Further, we addressed the question whether the immunologic profile represents an immune response indicative of prolonged or chronic antigenic exposure. Using machine learning, we identified algorithms that allowed for accurate determination of chronic COVID and severe COVID immunotypes. Further, we present a quantitative immunologic score that could be used to stratify patients to therapy and/or non-subjectively measure response to therapy.

## RESULTS

### Immune Profiling

To determine if immunologic abnormalities remain in Long Haulers, we performed high parameter immune cell quantification and characterization in a subset of individuals with preserved peripheral blood mononuclear cells. We determined B-cells, T-cells, and monocytes including subsets and including CD4/CD8 activation and exhaustion. Unlike active COVID-19, the CD4 and CD8 T-cell populations were within normal limits and there was no evidence of T-cell exhaustion (co-expression of PD-1, LAG3, and or CTLA-4). B-cells were significantly elevated compared to normal individuals (P<0.001) as was the CD14+, CD16+ monocytic subset (P<0.001) (Table 1). Interestingly, these two immune cell populations have been shown to be chronically infected by different viruses. B-cells are infected by Epstein-Barr and the CD14+, CD16+ monocytic subset by HIV-1 and by HCV (4).

**Table 1.** Immunologic parameters of study participants

| Average | CD3+% | CD4% | CD8% | CD4+PD1+% | CD4+LAG3+% | CD4+CTLA4+% | CD4+FoxP3+% | CD8+PD1+ % | CD8+LAG3+ % | CD8+CTLA4+% | CD19+% | CD14+CD16- % | CD16+CD14+% | CD16+CD14- % |
| --- | --- | --- | --- | --- | --- | --- | --- | --- | --- | --- | --- | --- | --- | --- |
| Normals | 64.45 | 53.89 | 33.83 | 35.62 | 0.94 | 1.51 | 6.21 | 43.76 | 4.35 | 1.39 | 6.04 | 42.79 | 9.00 | 32.68 |
| Lower CI | 54.39 | 43.21 | 27.20 | 28.36 | 0.49 | 0.75 | 4.54 | 33.50 | 2.71 | 0.74 | 5.04 | 34.41 | 4.60 | 25.49 |
| Upper CI | 74.50 | 64.57 | 40.46 | 42.89 | 1.39 | 2.26 | 7.87 | 54.01 | 5.99 | 2.03 | 7.04 | 51.16 | 13.41 | 39.86 |
| ong Hauler | 48.98 | 56.18 | 35.36 | 17.78 | 0.72 | 4.06 | 2.58 | 31.99 | 0.71 | 3.11 | 13.14 | 19.07 | 29.30 | 33.86 |

| Average<br>(pg/ml) | TNF-α | IL-4 | IL-13 | IL-2 | GM-CSF | sCD40L | CCL5<br>(RANTES) | CCL3<br>(MIP-1α) | IL-6 | IL-10 | IFN-γ | VEGF | IL-8 | CCL4<br>(MIP-1β) |
| --- | --- | --- | --- | --- | --- | --- | --- | --- | --- | --- | --- | --- | --- | --- |
| Normals | 9.09 | 4.18 | 3.94 | 6.17 | 51.27 | 7192.39 | 10781.84 | 22.82 | 2.21 | 0.67 | 1.94 | 9.32 | 16.87 | 76.84 |
| Lower CI | 7.37 | 2.17 | 1.79 | 5.53 | 25.72 | 5148.85 | 9764.99 | 13.05 | 1.65 | 0.42 | 0.63 | 6.36 | 13.03 | 61.00 |
| Upper CI | 10.81 | 6.18 | 6.09 | 6.82 | 76.82 | 9235.92 | 11798.68 | 32.60 | 2.77 | 0.92 | 3.26 | 12.28 | 20.72 | 92.67 |
| Long Haulers | 7.72 | 17.03 | 4.21 | 16.16 | 12.46 | 18302.41 | 12505.06 | 97.81 | 20.47 | 12.23 | 86.60 | 41.03 | 35.98 | 35.10 |
| Mild-Mod | 6.82 | 2.33 | 2.40 | 5.90 | 56.13 | 10673.72 | 11627.70 | 18.75 | 8.74 | 0.63 | 1.15 | 17.39 | 17.37 | 94.40 |
| Severe | 5.39 | 2.39 | 2.26 | 5.43 | 20.31 | 12306.39 | 11581.47 | 16.54 | 144.15 | 3.10 | 2.06 | 25.52 | 10.87 | 64.84 |

To further characterize the immune response in Long Haulers, we performed quantitative, multiplex cytokine/chemokine panel on 30 normal individuals to establish the normal range of the assay. We then analyzed 64 long haulers and compared the cytokine/chemokine profile (Table 1). IL-2, IL-4, CCL3, IL-6, IL-10, IFN-γ, and VEGF were all significantly elevated compared to normal control (all P<0.001). Conversely GM-CSF and CCL4 were significantly lower than normal controls. Further exacerbating this hyper-immunity was the significant decrease in T regulatory cells compared to normal individuals (P<0.001).

### Random Forest Binary and Multi-Class Models for Feature Selection and Prediction

We separated the dataset into a training and test split of 90% training and 10% test. This proportion was used because of the reduced number of instances in the dataset. Also, to ensure reproducible results we set the same random seed for all the models.

The first model we constructed was the multi-class predictor. This model attempted to separate the severe, long hauler and non-severe-non-long hauler class. This classifier achieved 97% precision, 97% recall and a F1 score of 0.97 in the training partition. In the test split, it performed slightly better, with a precision of 100%, a recall of 100% and thus and F1 score of 1.00 (Table 2). This model was then analyzed to identify the most relevant or informative features. This resulted in the identification of 6 features with an importance score above the importance median (0.063895) and average (0.07143). The identified features were: IFN-γ, IL-2, IL-6, IL-10, IL-8, CCL4-MIP-1β, in importance order. The full list of ranked features can be seen in figure 2.

**Figure 2.**
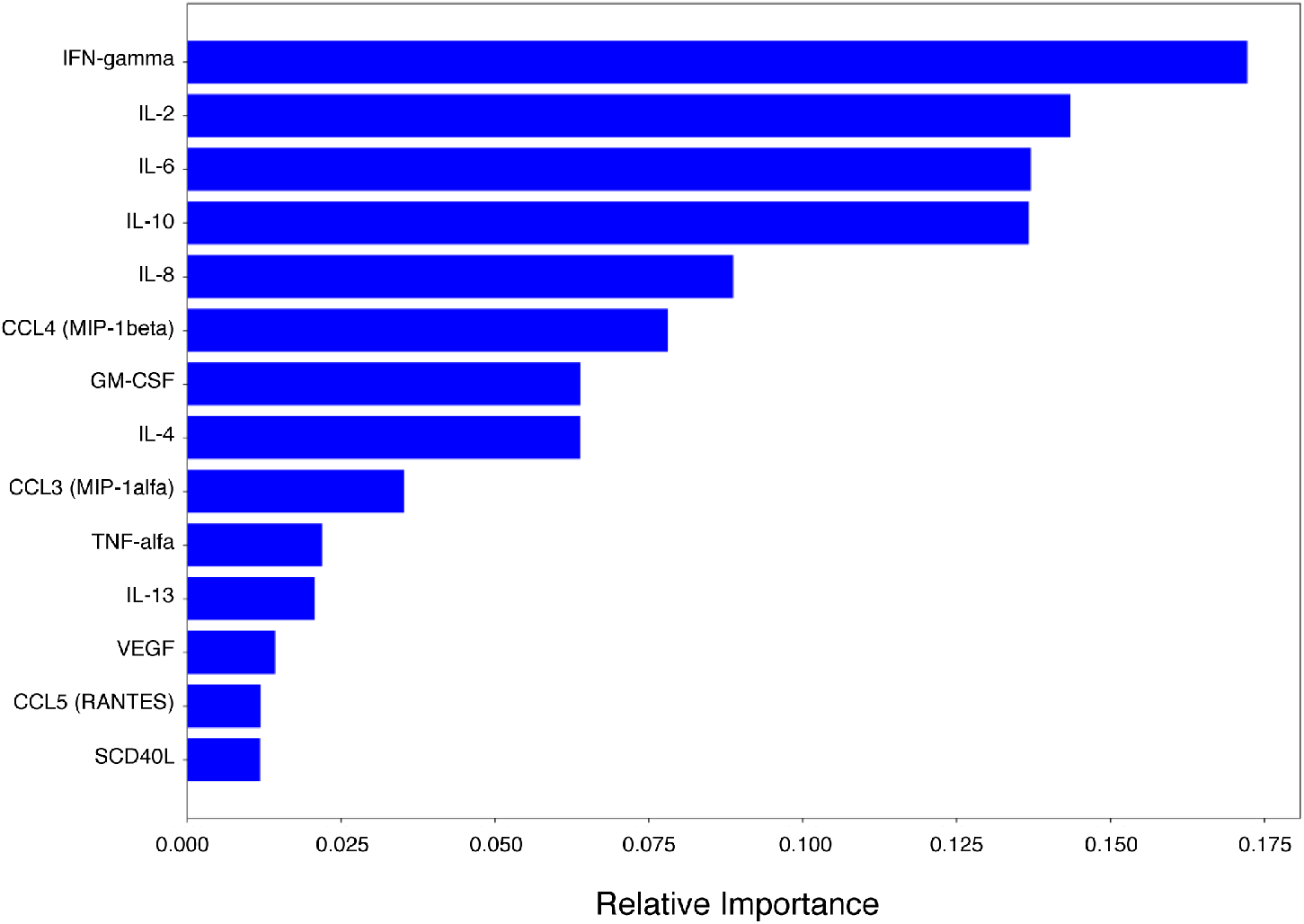
Feature importance for multi-class classifier using Random Forest predictor.

**Table 2.** Performance Metrics for the Random Forest Classifiers in the test split.

| Model | Precision % | Recall % | F1 Score |
| --- | --- | --- | --- |
| Long Hauler-Full Features | 100 | 100 | 1.00 |
| Severe- Full Features | 100 | 100 | 1.00 |
| Multi-Class-Full Features | 100 | 100 | 1.00 |

Regarding the long hauler and non-long hauler binary classifier, our results were consistent between the training and the test set. In both partitions the precision and the recall were 100% (1.00) and thus the F1 score equaled 1.00. The observation that the model had good metrics in the test split when compared to the train set is a valuable indicator that the model is not overfitting, and that it is capable of generalizing the patters identified in the training data. The overview of the precision, recall and F1 score for the binary long hauler model can be seen in table 2. Feature importance analysis of the binary model, revealed that the features identified as important for this model were the same features identified as important for the multi-class predictor. This finding suggests there is an important group of characteristics or variables that are influential in the identification of long hauler data points from other instances. These features can be seen in figure 2.

The severe binary model, which classified instances between non-severe and severe resulted in high performance metrics for both the training and test splits. As shown in table 2, the performance of this model was an indicator of no potential overfitting. This model is of special interest given the small number of instances in the severe class. Furthermore, the feature importance analysis of this model revealed that the relevant features were also the same as with the multi-class model and with the long hauler binary classifier (Figure 2). This finding reinforces our notion that these group of relevant features could impact classification, or that could have some biological significance worth exploring by means of other analysis like a separation heuristic.

### Deep Neural Network Binary Classifiers using the Full Feature Set

The deep neural network (DNN) classifier was constructed layers of neurons. Each layer transformed the inputs inputs using the rectified linear activation function or ReLU. The DNN model was constructed to have 1 input layer, 3 hidden layers with 10 neurons each, followed by layer with 6 neurons. Finally, the output layer consists of 3 neuros, for the outputs (classes) and the softmax (multi-class) or sigmoid (binary) function. This architecture was used for the multi-class model and the binary models.

The results of the long hauler binary models, revealed differences of ~5% between the metrics of the training and the test set (Table 3). Such difference is not significant to attribute overfitting to the training set. In contrast, the severe binary model had significant differences between the performance metrics of the training and the test set (Table 3). This is evident in the precision score, with 98% in the training set and 75% on the test set, and thus the F1 score with a difference of 20% (0.99 on the training set and 0.79 on the test set). A potential explanation could be that the severe class has a limited number of data points, but our random forest classifier for the severe class perfumed well. These results suggest that the best approach is a multi-class predictor.

**Table 3.** Performance Metrics of the DNN full feature model in the training and test splits

| DNN | Precision % | Recall % | F1 Score |
| --- | --- | --- | --- |
| Multi-Class -<br>Full Features<br>-Train | 99 | 97 | 0.98 |
| Long Hauler -<br>Full Features<br>-Train | 100 | 100 | 1.00 |
| Severe - Full<br>Features -<br>Train | 98 | 100 | 0.99 |
| Multi-Class -<br>Full Features<br>-Test | 100 | 100 | 1.00 |
| Long Hauler -<br>Full Features<br>-Test | 94 | 94 | 0.93 |
| Severe - Full<br>Features -<br>Test | 75 | 92 | 0.79 |

### Multi-class Deep Neural Network Classifiers using the Full Feature Set

The multi-class DNN implemented using the full feature set had good metrics (Table 3). The precision, recall and F1 score of 100%, 100% and 1.00 in the test split. This indicates that the model is not overfitting, and validating our notion that this would generalize better than the binary models. The model’s performance is supported by its confusion matrix (true class vs predicted) where it is possible to determine how well it can predict the three classes (Figure 3).

**Figure 3.**
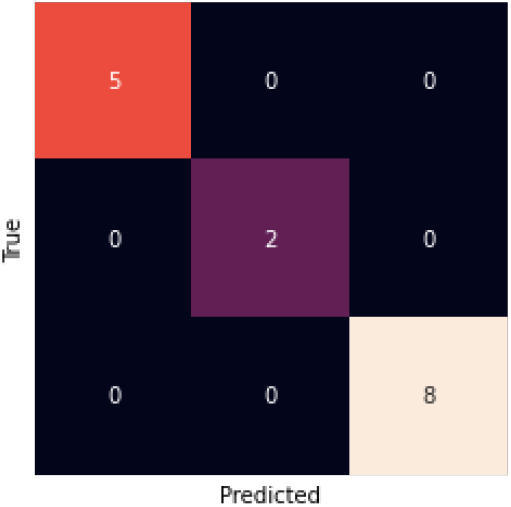
Full-feature multi-class DNN model confusion matrix for the test split.

The potential of a DNN classifier is that it adjusts multiple parameters transform the inputs into outputs. This is very important because the vast number of parameters allows for the model to better identify hidden signals in the data. Also, DNN require hyperparameter tuning, such as learning rate, number of hidden layers and neurons per hidden layer, as well as the optimizer and activation function, which affect the performance of the model. By adjusting these hyperparameters and castrating a model capable of finding the hidden relationships in the data we were able to achieve such high results and construct a predictive multi-class system.

### Reduced Feature Multi-class Deep Neural Network Classifiers

The results of the DNN indicated that the multi-class had the highest performance. Based on this, we constructed a DNN using the 6 most important features identified by the random forest variable importance. This model was known as minimal DNN or mDNN. This model was constructed using the same architecture as the full feature set DNN. This model’s performance in the training set and the test set (Table 4), revealed a significant difference in both precision and recall, such difference could indicate that although the 6 features were identified as the most relevant, it could be possible that all variables contribute to the hidden pattern that makes up the classification of the instances. This idea is supported by the differences in performance between the mDNN and the full feature classifier in both training and test splits (Tables 3 & 4). This is further supported by the comparison of the confusion matrices, where mDNN (figure 4A) misclassifies more instances than the full feature multi-class DNN (Figure 3).

**Figure 4.**
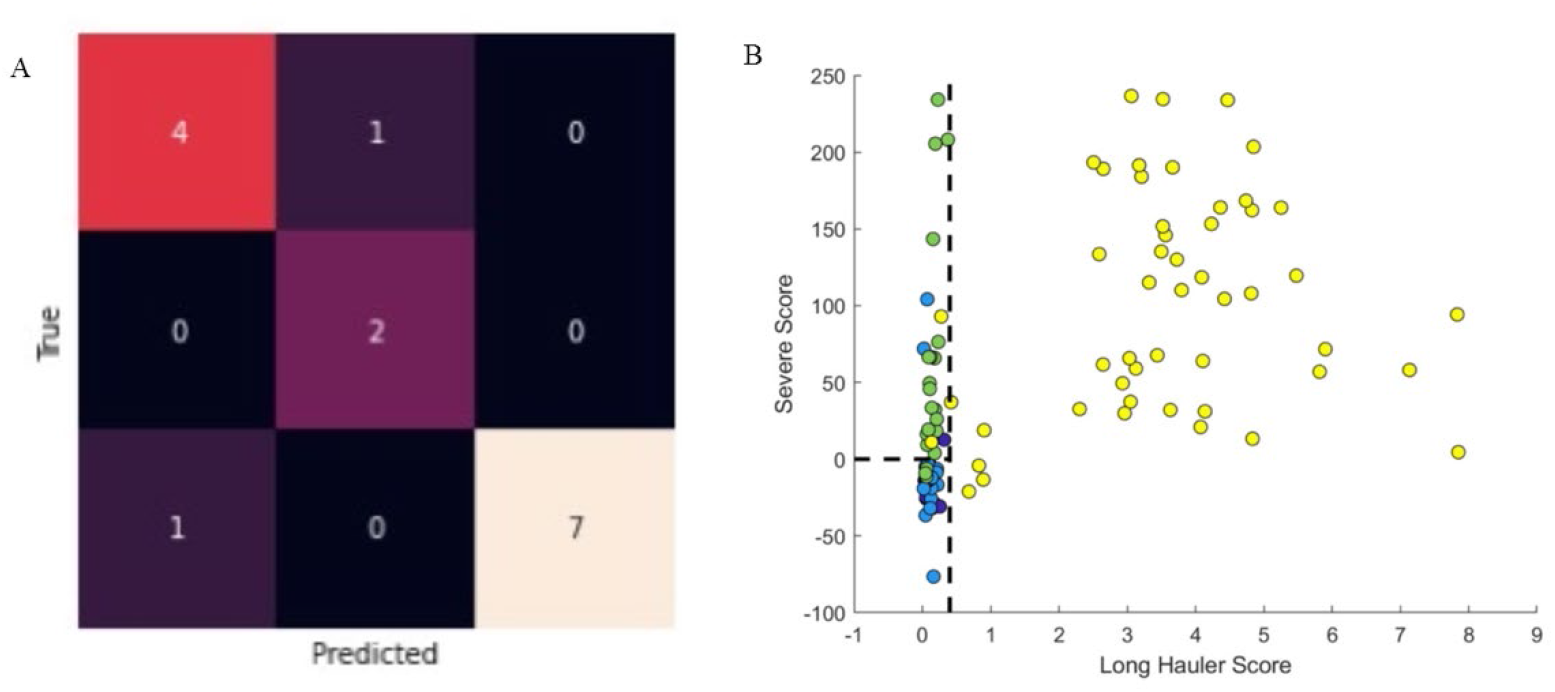
Classification abilities of the minimal Deep Neural Network (mDNN) and the discrimination heuristic generated using important variables. A) The confusion matrix for the mDNN classifier denoting the presence of false positives for the severe and other classes. B) Discrimination ability of the heuristic with reduced or most important features identified using Random Forest classifier. The dots represent the data points, where yellow are long haulers, green-severe, dark blue-mild/moderate and light blue-normal.

**Table 4.**
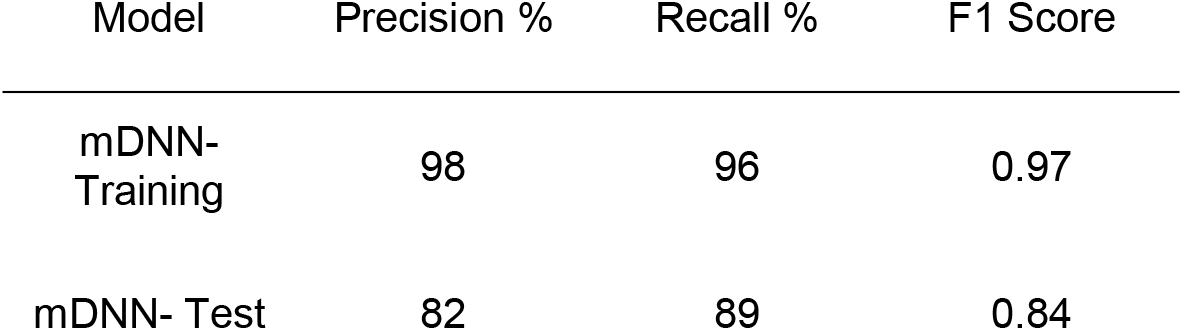
Performance metrics for the minimal deep neural network (mDNN) on the training and test splits.

Moreover, we simplified our prediction model by feature engineering of two classification scores based on the top informative features. First, a “Long Hauler Score” was defined as S1 = (IFN-γ + IL-2) / CCL4-MIP-1β. Second, “Severe Score” was defined as S2 = (10*IL-10 + IL-6) – (IL-2 + IL8). Using a combined heuristic to first classify the Long Haulers (S1>0.4) and second the severe COVID-19 patients (S2>0), we obtained a sensitivity of 97% for Long Haulers with a 100% specificity and a sensitivity of 88% for severe patients with a specificity of 96% (Figure 4B).

## DISCUSSION

Individuals infected with SARS-Cov2 exert distinct severity patterns which have been associated with different immune activation profiles. Interestingly, in some cases longer times are required to experience full recovery, representing a particular pathological type recently described as long-COVID or long haulers (LH). The scientific evidence generated during the last months strongly supports that the different outcomes on COVID-19 patients are determined by the immune mechanisms activated in response to the viral infection.

The immune response to SARS-Cov2 induces a release of different molecules with inflammatory properties such as cytokines and chemokines. This event, known as cytokine storm, is an immunopathological feature of COVID-19 and it has been associated with the severity of the disease. The increase in blood concentrations of different cytokines and chemokines such as IL-6, IL-8, IL-10, TNF-α, IL-1β, IL-2, IP-10, MCP-1, CCL3, CCL4, and CCL5 has been described for COVID-19 patients (5). Some of these molecules have been proposed as biomarkers to monitor the clinical evolution and to determine treatment selection for COVID-19 patients. Nevertheless, it is important to consider that some of these molecules function in a context dependent manner, therefore the clinical relevance of analyzing single cytokine changes is limited.

One of the most important challenges during the pandemics is to avoid the saturation of the health systems, therefore the determination of predictive biomarkers that allow a better stratification of the patients is paramount. Even though cytokines such as IL-6 and IL-8 have been proposed as indicators of the disease severity, and in some studies they were strong and independent predictors of patient survival (6), their predictive value when analyzed alone is debatable (7). The generation of scores considering blood levels of cytokines and chemokines with different immunological functions incorporates the importance of the context-dependent function of these molecules.

In order to predict severe cases, a score was generated considering IL-10, IL-6, IL-2, and IL-8 blood concentrations. In this classification, severe cases are characterized by high IL-6 and IL-10 levels, both cytokines previously attributed to increase the immunopathogenesis of COVID-19 and predictive value in severe cases (6, 8). In different settings, IL-6 has been associated with oxidative stress, inflammation, endothelial dysfunction, and thrombogenesis (9-12) which are characteristic features of severe COVID-19 cases caused by excessive myeloid cell activation (13). Consistently, increased IL-10 levels interfere with appropriate T-cell responses, inducing T-cell exhaustion and regulatory T cell polarization leading to an evasion of the anti-viral immune response (14). Furthermore, besides its anti-inflammatory function on T cells, in some settings IL-10 induces STAT1 activation and a pro-inflammatory response in type I IFN-primed myeloid cells (15,16). Therefore, elevated levels of IL-6 and IL-10 promote myeloid cell activation, oxidative stress, endothelial damage, and dampens adequate T cell activation. Additionally, to strengthen the classification, the score presented here, differentiates the severe cases by the subtraction of IL-2 and IL-8, which are cytokines related to proper T cell activation (IL-2) and recruitment (IL-8).

According to the score generated for distinguishing LH, these patients are characterized by an increased IFN-γ and IL-2 and a reduced CCL4 production. In the context of a viral infection, the combination of IFN-γ and IL-2 would induce the activation of effector T cells with pro-inflammatory properties and the capacity of generating an effective immune response to eliminate the virus. However, LH are characterized by longer periods of time with clinical signs and symptoms such as fatigue and lung damage. This suggests that the inflammatory context created by these cytokines to induce T cell activation is not enough to generate an adequate anti-viral response without the proper recruitment signals to attract activated T cells. CCL4 signals through the receptor CCR5 to attract T cells to the site of inflammation and depending on the immune context, this molecule recruits differently activated T cells (17,18). Moreover, it was recently shown by single cell analysis a down regulation of CCL4 expression in peripheral myeloid cell compartments in patients with mild and severe COVID-19 (19). In LH, IFN-γ and IL-2 would create an immune context to induce Th1 polarization, but the low levels of CCL4 affect the recruitment of these cells impairing the antiviral response. The effect of increased IFN-γ and IL-2 on T cell activation is evident in the reduction of the percentage of exhausted (CD4+PD1+/ CD8+PD1+) and regulatory T cells (FoxP3+) compared to healthy donors. Interestingly, there is an increase in the percentage of circulating CD4+ and CD8+ T cells expressing CTLA-4 in the LH group compared to healthy donors, which is a molecule that affects antigen presentation in secondary lymphoid organs, but its presence in circulating T cells may reflect a compensatory mechanisms to the low CCL4 levels in the LH group. CTLA-4 induced signaling in T cells upregulates the expression of the CCL4 receptor CCR5 (20, 21), in the LH group CTLA-4 upregulation suggests a failed attempt to increase the sensitivity of IFN-γ/IL-2 activated T cells to CCL4. Therefore, proper T cell activation (high IFN-γ+IL-2) but ineffective T cell recruitment (low CCL4) are characteristic features of the failed antiviral response observed in the LH group supporting virus persistence. Additionally, increased IFN-γ promotes myeloid cell activation which is observed in the augmented percentage of inflammatory CD14+CD16+ monocytes in the LH group compared to healthy donors, supporting lymphopenia and virus persistence in these patients. This is supported by recent findings describing an increased gene expression in response to IFN-γ in mild and severe COVID-19 patients in peripheral myeloid cells (19) and the dysregulation in the balance of monocyte populations by the expansion of the monocyte subsets described in COVID-19 patients (22). Finally, we propose that long-lasting pulmonary damage observed in LH, is caused by a combination of factors including 1) longer virus persistence influenced by LH immune profile characterized by high IFN-γ and IL-2 levels inducing Th1 polarization which is ineffective with low CCL4-induced T cell recruitment, leading to an inflammatory myeloid cell activation; and 2) the immunopathological pulmonary effects consequence of this LH immune profile.

Regarding the immunopathological effects of LH immune profile, using murine models it has been shown that high IFN-γ levels could affect the kinetics of the resolution of inflammation-induced lung injury as well as thrombus resolution (23, 24), which could be related to long-lasting symptoms of LH associated to pulmonary coagulopathy and immune-mediated tissue damage.

Interestingly, COVID-19 individuals (including LH, mild, severe) show high levels of CCL5, a chemoattractant that like CCL4 signals through CCR5. Indeed, the disruption of the CCL5-CCR5 pathway restores immune balance in critical COVID-19 patients (4). In the specific case of LH, despite the high concentrations of CCL5 a reduction on the CCL4-mediated recruitment of activated T cells is proposed. This could be related to different factors:

1. Reduction of total recruitment signals in LH with low CCL4 concentrations.
2. Different functional responses of CCL4 and CCL5 to polymorphic variants of the CCR5. Distinct functional features have been reported to CCR5 variants regarding binding avidity, receptor internalization, Ca++ influx and chemotactic activity (25). Even though, clear mechanistic differences between CCL4 and CCL5 interaction with CCR5 are missing, it has been suggested that is important to consider the knowledge gained on CCR5 polymorphisms in HIV/AIDS context (26).
3. Signaling through alternative receptors for CCL5. Besides CCR5, CCL5 can signal through the receptors CCR1 and CCR3 (27) whereas CCL4 effects are restricted to CCL5. It has been shown that CCL4 can bind to CCR1 but is not able to induce the intracellular pathway necessary for activating the chemoattractant stimulus (27,28). Therefore, CCL4 has been proposed as an antagonist of CCR1 (28), however further analysis of this needs to be performed. Interestingly, CCR1 is expressed on blood myeloid cells such as monocytes and neutrophils (27), and it is upregulated on COVID-19 patients (29). Additionally, high levels of IFN-γ (feature of LH) have been associated with an increase CCR1 expression on human neutrophils (30). Therefore, in LH, high levels of CCL5 (combined with low levels of potential CCR1-antagonist CCL4) leads to a higher recruitment of myeloid cells expressing CCR1.

## MATERIAL/METHODS

### Patients

Following informed consent, whole blood was collected in a 10 mL EDTA tube and a 10 mL plasma preparation tube (PPT). A total of 144 individuals were enrolled in the study consisting of 29 normal individuals, 26 mild-moderate COVID-19 patients, 25 severe COVID-19 patients and 64 chronic COVID (long hauler-LH) individuals. Long Haulers symptoms are listed in Figure 1. Study subjects were stratified according to the following criteria.

**Figure 1.**
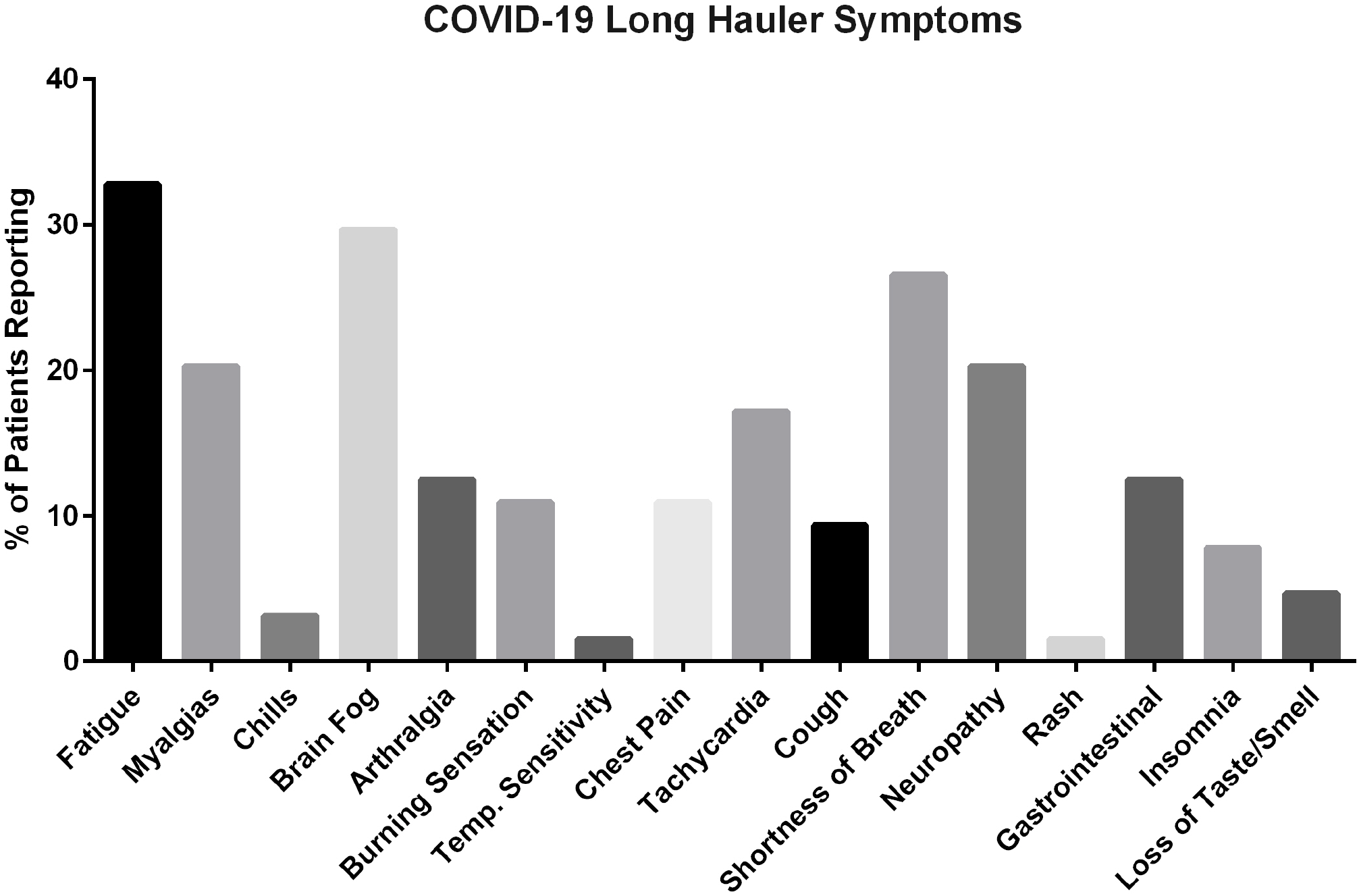
Symptoms reported by long hauler patients enrolled in the study.

#### Mild

1. Fever, cough, sore throat, malaise, headache, myalgia, nausea, diarrhea, loss of taste and small
2. No sign of pneumonia on chest imaging (CXR or CT Chest)
3. No shortness of breath or dyspnea

#### Moderate

1. Radiological findings of pneumonia fever and respiratory symptoms
2. Saturation of oxygen (SpO2) ≥ 94% on room air at sea level

#### Severe

1. Saturation of oxygen (SpO2) < 94% on room air at sea level
2. Arterial partial pressure of oxygen (PaO2)/ fraction of inspired oxygen (FiO2) < 300mmHG
3. Lung infiltrate > 50% within 24 to 48 hours
4. HR ≥ 125 bpm
5. Respiratory rate ≥ 30 breaths per minute

#### Critical

1. Respiratory failure and requiring mechanical ventilation, ECMO, high-flow nasal cannula oxygen supplementation, noninvasive positive pressure ventilation (BiPAP, CPAP)
2. Septic Shock-Systolic blood pressure < 90mmHg or Diastolic blood pressure< 60 mmHg or requiring vasopressors (levophed, vasopressin, epinephrine
3. Multiple organ dysfunction (cardiac, hepatic, renal, CNS, thrombotic disease)

#### Post-acute COVID-19 (Long COVID)

1. Extending beyond 3 weeks from the initial onset of first symptoms

#### Chronic COVID-19

1. Extending beyond 12 weeks from the initial onset of first symptoms (Table 1)

### High Parameter Immune Profiling/Flow Cytometry

Peripheral blood mononuclear cells were isolated from peripheral blood using Lymphoprep density gradient (STEMCELL Technologies, Vancouver, Canada). Aliquots 200 of cells were frozen in media that contained 90% fetal bovine serum (HyClone, Logan, UT) and 10% dimethyl sulfoxide (Sigma-Aldrich, St. Louis, MO) and stored at - 70°C. Cells were stained and analyzed as previously described (4) (Patterson) using a 17-color antibody cocktail.

### Multiplex Cytokine Quantification

Fresh plasma was used for cytokine quantification using a customized 14-plex bead based flow cytometric assay (IncellKINE, IncellDx, Inc) on a CytoFlex flow cytometer as previously described using the following analytes: ‘TNF-α’, ‘IL-4’, ‘IL-13’,’IL-2’, ‘GM-CSF’, ‘sCD40L’, ‘CCL5 (RANTES)’, ‘CCL3 (MIP-1α)’,’IL-6’, ‘IL-10’, ‘IFN-γ’, ‘VEGF’, ‘IL-8’, and ‘CCL4 (MIP-1β) (4). For each patient sample, 25 μL of plasma was used in each well of a 96-well plate. Standard curves with serial 6 point dilutions of antigen were run on each plate for each cytokine. Raw data was analyzed using LegendPlex software (Biolegend, Inc San Diego CA). Samples were run in duplicate.

### Data Processing

Data was imported and processed using Python 2.7, using the *pandas* library (version 1.1.0). and the numeric python module, *numpy* version 1.18.5. Our data consisted of 144 instances representing 4 classes (Normal-n=29, Mild-Moderate-n=26, Severe-n=25, Long Hauler-n=64). Each class had 14 columns, representing the different cytokine/chemokine analytes. Each analyte had different measurements which required a normalization process to reduce outlier effect and to facilitate algorithm convergence.

Normalization was done using Min-Max and based on a linear transformation of the original data. Min-Max maintains the original relationship between the data, while fitting it within a pre-defined boundary. The Python implementation of min-max calculates the range in such a manner that the range of the features will be defined between 0 and 1. For this reason, min-max normalization is also referred to as 0-1 normalization (or scaling). The typical min-max transformation is given in equation 1:

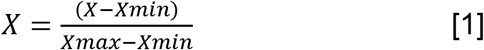

### Target Variable Processing

Since Min-max normalization, can only be applied to numeric variables a new variable defined as *targets* was created.The variable targets represent the different classes (Long Hauler, Severe, Mild-Moderate, and Normal) for the instances in the dataset. The resulting array has 4 classes for each state. The goal of our analysis is to properly identify/discriminate the instances that belong to the Severe state or the Long-Hauler state compared to other states. This goal can be achieved by building either binary classifiers for the Severe class and for the Long Hauler class, a multi-class predictor. For the construction of both models, t is required to separate the targets to reflect the dosing question: can a predictor discriminate between the Severe, Long Hauler and Other Sates.

To build the models that answer this question, we grouped the M-M and Normal labels in a new class which was distinct form the Severe and Long-Hauler states. We then proceeded to apply filters based on the task (binary or multi-class classification). For the Severe binary predictor, we conditioned the targets to be exactly Severe or else they were assigned to Not-Severe. This same task was done for Long-Haulers, were either an instance label was exactly labelled Long-Hauler or else it would be assigned to the Non-Long Hauler class. The multi-class predictor processing only requires to define three classes: Severe, Long-Hauler and Non-Severe-Non-Long-Hauler which was composed of the Normal and Mild-Moderate cases.

### One-hot Encoding of Targets

The implementation of one-hot encoding on the target variable, is based on the notion that multiple machine learning algorithms are unable to properly process categorical data. It is possible to use numeric replacements, such as integer values, but this can only be useful if there is an ordinal relationship within the variable. Such use would imply that there exists a vectorial relationship between the labels, for example, in our classes we have Normal, Mild-Moderate, Severe and Long-Haulers. If we assigned a vector of integers from 0 to 4 in their corresponding orders to the classes, it would assume the presence of a vectorial distance between Normal and Long Hauler or V0 –> V4.

To properly design an experiment that reflects this, we use one-hot encoding After applying one-hot encoding the labels are substituted with 1 and 0, where 1 represents the presence of the class and 0 the absence. The use of one-jot encoding corrects for the vector-distance assumption of integer or categorical classes, where higher or larger values could be interpreted as better.

Definition of precision, recall and F1 score

The precision (equation 2) is a measure of the percentage of the results that are relevant. The metric Recall measures the percentage of the total relevant results that are correctly classified by the predictor (equation 3). The harmonic mean between these two measures is known as the F1 score and ranges from 0 to 1, the closer to is to 1, the better the model performs (equation 4). The F1 score for both false positives (FP) and false negatives (FN) as well as for true positives (TP).

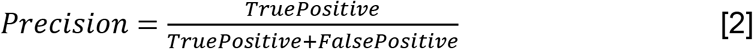

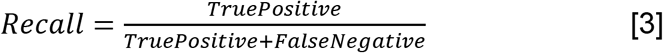

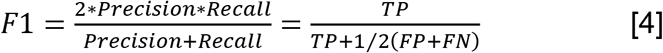

### Feature Selection and Classification using Random Forest

Data pre-processing, target variable processing and the encoding of targets were performed before classification as above. Feature selection is the process of reducing dimensionality of the dataset by selecting those features or variables that are more informative than those that are not.

To perform feature selection, we implemented the RandomForestClassifier method from Sci-kit Learn. Random Forest allows for identification of features that better separate the classes by determining what percentage of the nodes that use those features have a reduction in entropy or impurity (which are measures of how well separated the instances are using a feature).

The binary classifier was constructed using the data points and their features with the one-hot encoded target corresponding to: 1) the severe and non-severe model, 2) the long hauler and non-long hauler model and 3) the multiclass model. The model was built with the RandomForestClassifier method from Sci-kit Learn, with the number of trees constructed set to 750, the number of features set as the square root of the feature space, and the node depth equal to 4 to avoid overfitting. These parameters were set for binary and multi-class predictors. Model performance was measured using: precision, recall and the F1 score (see supplementary information).

### Predictor Construction Using Deep Neural Networks

The deep neural network (DNN) binary and multiclass classifiers were constructed with a basic DNN architecture built on stacks of perceptrons, where each subsequent layer is connected to the previous one. Each layer transformed the inputs inputs using the rectified linear activation function or ReLU. The DNN models were constructed to have 1 input layer, 3 hidden layers with 10 neurons each, followed by layer with 6 neurons.

Finally, the output layer consists of 3 neurons, for the outputs (classes) and the softmax (multi-class) or sigmoid (binary) function.

In order for a DNN to generate the best possible predictions, we minimized the loss function or error of the model using the ADAM optimizer to search for the optimal combination of hyperparameters. When setting the optimizer, we defined the learning rate to 1e-3. The loss function was set to categorical cross entropy because the targets are one-hot encoded.

## Acknowledgments

The authors would like to acknowledge the work of Christine Meda in coordinating the study and interacting with the patients,

## Author contributions

R.Y. organized the clinical study and actively recruited patients.

B.K.P, A.P., H.R., E.L. performed experiments and analyzed the data.

J.G-C., R.A.M., J.M. performed the bioinformatics

B.K.P., J.M., J.G-C., R.A.M. wrote the draft of the manuscript and all authors contributed to revising the manuscript prior to submission.

## Data and materials availability

All requests for materials and data should be addressed to the corresponding author

## Abbreviations

IL: interleukin
RANTES: regulation on activation
normal T: expressed and secreted
CCR: chemokine receptor
IFN: interferon
TNF: tumor necrosis factor
MIP: macrophage inflammatory protein
GM-CSF: granulocyte-macrophage colonystimulating factor
VEGF: vascular endothelial growth factor
HIV: human immunodeficiency virus
HCV: hepatitis C virus

## REFERENCES

1. L. Chen, H. Deng, H. Cui, J. Fang, Z. Zuo, J. Deng, Y. Li, X. Wang, L. Zhao Inflammatory responses and inflammation-associated diseases in organs. Oncotarget 9, 7204–7218 (2018).

2. S. Rasa, Z. Nora-Krukle, N. Henning et al. Chronic viral infections in myalgic encephalomyelitis/chronic fatigue syndrome (ME/CFS). J Transl Med 16, 268 (2018).

3. P.A. Mudd, J.S. Turner, A. Day, W.B. Alsoussi, Z. Liu, J.A. O’Halloran, R.M. Presti, B.K. Patterson, S.P.J. Whelen, A. Ellebedy. SARS-CoV-2 viral RNA shedding for more than 87 days in an individual with an impaired CD8+ T-cell response. Front Immunol (in press).

4. G. Coquillard, B. Patterson. HCV-Infected, Monocyte Lineage Reservoirs Differ in Individuals with or without HIV Co-Infection. J Infect Dis 2009;200:947–954.

5. B. K. Patterson, H. Seethamraju, K. Dhody, M. J. Corley, K. Kazempour, J. P., Lalezari, A. P. Pang, C. Sugai, E. B. Francisco, A. Pise, H. Rodrigues, M. Ryou, H. L. Wu, G. M. Webb, B. S. Park, S. Kelly, N. Pourhassan, A. Lelic, L. Kdouh, M. Herrera, E. Hall, E. Aklin, L. Ndhlovu, J. B. Sacha. CCR5 inhibition in Critical COVID-19 Patients Decreases Inflammatory Cytokines, Increases CD8 T-Cells, and Decreases SARS-CoV2 RNA in Plasma by Day 14. Int J Infect Dis (2020) doi:10.1016/j.ijid.2020.10.101

6. D. M. Del Valle, S. Kim-Schulze, H. H. Huang, N. D. Beckmann, S. Nirenberg, B. Wang, Y. Lavin, T. H. Swartz, D. Madduri, A. Stock, T. U. Marron, H. Xie, M. Patel, K. Tuballes, O. Van Oekelen, A. Rahman, P. Kovatch, J. A. Aberg, E. Schadt, S. Jagannath, M. Mazumdar, A. W. Charney, A. Firpo-Bncourt, D. R. Mendu, J. Jhang, D. Reich, K. Sigel, C. Cordon-Cardo, M. Feldmann, S. Parekh, M. Merad, S. Gnjatic,. An inflammatory cytokine signature predicts COVID-19 severity and survival. Nat Med, 26, 1636–1643 (2020).

7. S. M. Russell, A. Alba-Patiño, E. Barón, M. Borges, M. Gonzalez-Freire, & R. de la Rica. Biosensors for Managing the COVID-19 Cytokine Storm: Challenges Ahead. ACS Sens, 5, 1506–1513.

8. S. K. Dhar, K, V., S. Damodar, S. Gujar & M. Das, IL-6 and IL-10 as redictors of disease severity in COVID 19 patients: Results from Meta-analysis and Regression. medRxiv, (2020). 2008.2015.20175844. https://doi.org/10.1101/2020.08.15.20175844

9. T. Hou, Tieu, B. C., S. Ray, A. Recinos Iii, R. Cui, R. G. Tilton, & A. R. Brasier. Roles of IL-6-gp130 Signaling in Vascular Inflammation. Curr Cardiol Rev, 4, 179–192. https://doi.org/10.2174/157340308785160570

10. J. Lee, S. Lee, H. Zhang, M. A. Hill, C. Zhang, & Y. Park. Interaction of IL-6 and TNF-α contributes to endothelial dysfunction in type 2 diabetic mouse hearts. PLoS One, 12, e0187189. https://doi.org/10.1371/journal.pone.0187189

11. V. Roldán, F. Marín, A. D. Blann, A. García, P. Marco, F. Sogorb, & G. Y. Lip. Interleukin-6, endothelial activation and thrombogenesis in chronic atrial fibrillation. Eur Heart J, 24, 1373–1380. https://doi.org/10.1016/s0195-668x(03)00239-2

12. S. Wassmann, M. Stumpf, K. Strehlow, A. Schmid, B. Schieffer, M. Böhm, & G. Nickenig. Interleukin-6 induces oxidative stress and endothelial dysfunction by overexpression of the angiotensin II type 1 receptor. Circ Res, 94, 534–541. https://doi.org/10.1161/01.res.0000115557.25127.8d

13. D. McGonagle, K. Sharif, A. O’Regan, & C. Bridgewood. The Role of Cytokines including Interleukin-6 in COVID-19 induced Pneumonia and Macrophage Activation Syndrome-Like Disease. Autoimmun Rev, 19, 102537. https://doi.org/10.1016/j.autrev.2020.102537

14. J. M. Rojas, M. Avia, V. Martín, & N. Sevilla. IL-10: A Multifunctional Cytokine in Viral Infections. J Immunol Res, 2017, 6104054. https://doi.org/10.1155/2017/6104054

15. H. Mühl. Pro-Inflammatory Signaling by IL-10 and IL-22: Bad Habit Stirred Up by Interferons? Front Immunol, 4, 18. https://doi.org/10.3389/fimmu.2013.00018

16. M. N. Sharif, I. Tassiulas, Y. Hu, I. Mecklenbräuker, A. Tarakhovsky, & L. B. Ivashkiv. IFN-alpha priming results in a gain of proinflammatory function by IL-10: implications for systemic lupus erythematosus pathogenesis. J Immunol, 172, 6476–6481. https://doi.org/10.4049/jimmunol.172.10.6476

17. J. Y. Liu, F. Li, L. P. Wang, X. F. Chen, D. Wang, L. Cao, Y. Ping, S. Zhao, B. Li, S. H. Thorne, B. Zhang, P. Kalinski, & Y. Zhang. CTL-vs Treg lymphocyte-attracting chemokines, CCL4 and CCL20, are strong reciprocal predictive markers for survival of patients with oesophageal squamous cell carcinoma. Br J Cancer, 113, 747–755. https://doi.org/10.1038/bjc.2015.290

18. N. Mukaida, S. I. Sasaki, & T. Baba. CCL4 Signaling in the Tumor Microenvironment. Adv Exp Med Biol, 1231, 23–32. https://doi.org/10.1007/978-3-030-36667-43

19. G. Xu, F. Qi, H. Li, Q. Yang, H. Wang, X. Wang, X. Liu, J. Zhao, X. Liao, Y. Liu, L. Liu, S. Zhang, & Z. Zhang. The differential immune responses to COVID-19 in peripheral and lung revealed by single-cell RNA sequencing. Cell Discov, 6, 73. https://doi.org/10.1038/s41421-020-00225-2

20. K. Knieke, H. Hoff, F. Maszyna, P. Kolar, A. Schrage, A. Hamann, G. F. Debes, M. C. Brunner-Weinzierl.. CD152 (CTLA-4) determines CD4 T cell migration in vitro and in vivo. PLoS One, 4, e5702. (2009). https://doi.org/10.1371/journal.pone.0005702

21. K. Knieke, H. Lingel, K. Chamaon, & M. C. Brunner-Weinzierl. Migration of Th1 lymphocytes is regulated by CD152 (CTLA-4)-mediated signaling via PI3 kinase-dependent Akt activation. PLoS One, 7, e31391. (2012). https://doi.org/10.1371/journal.pone.0031391

22. W. Shi, X. Liu, Q. Cao, P. Ma, W. Le, L. Xie, J. Ye, W. Wen, H. Tang, W. Su, Y. Zheng, & Y. Liu. High-dimensional single-cell analysis reveals the immune characteristics of COVID-19. Am J Physiol Lung Cell Mol Physiol. https://doi.org/10.1152/ajplung.00355.2020

23. J. R. Mock, M. K. Tune, C. F. Dial, J. Torres-Castillo, R. S. Hagan, & C. M. Doerschuk. Effects of IFN-γ on immune cell kinetics during the resolution of acute lung injury. Physiol Rep, 8, e14368. https://doi.org/10.14814/phy2.14368

24. M. Nosaka, Y. Ishida, A. Kimura, Y. Kuninaka, M. Inui, N. Mukaida, & T. Kondo. Absence of IFN-γ accelerates thrombus resolution through enhanced MMP-9 and VEGF expression in mice. J Clin Invest, 121, 2911–2920. https://doi.org/10.1172/jci40782

25. H. F. Dong, K. Wigmore, M. N. Carrington, M. Dean, J. A. Turpin, & O. M. Howard. Variants of CCR5, which are permissive for HIV-1 infection, show distinct functional responses to CCL3, CCL4 and CCL5. Genes Immun, 6, 609–619. https://doi.org/10.1038/sj.gene.6364247

26. R. K. Mehlotra. Chemokine receptor gene polymorphisms and COVID-19: Could knowledge gained from HIV/AIDS be important? Infect Genet Evol, 85, 104512. https://doi.org/10.1016/j.meegid.2020.104512

27. C. E. Hughes, & R. J. B. Nibbs. A guide to chemokines and their receptors. Febs j, 285, 2944–2971. https://doi.org/10.1111/febs.14466

28. H. Gaertner, O. Lebeau, I. Borlat, F. Cerini, B. Dufour, G. Kuenzi, A. Melotti, R. J. Fish, R. Offord, J. Y. Springael, M. Parmentier, & O. Hartley. Highly potent HIV inhibition: engineering a key anti-HIV structure from PSC-RANTES into MIP-1 β/CCL4. Protein Eng Des Sel, 21, 65–72. https://doi.org/10.1093/protein/gzm079

29. P. R. Ray, A. Wangzhou, N. Ghneim, M. S. Yousuf, C. Paige, D. Tavares-Ferreira, J. M. Mwirigi, S. Shiers, I. Sankaranarayanan, A. J. McFarland, S. V. Neerukonda, S. Davidson, G. Dussor, M. D. Burton, & T. J. Price. A pharmacological interactome between COVID-19 patient samples and human sensory neurons reveals potential drivers of neurogenic pulmonary dysfunction. Brain Behav Immun, 89, 559–568. https://doi.org/10.1016/j.bbi.2020.05.078

30. R. Bonecchi, N. Polentarutti, W. Luini, A. Borsatti, S. Bernasconi, M. Locati, C. Power, A. Proudfoot, T. N. Wells, C. Mackay, A. Mantovani, & S. Sozzani. Up-regulation of CCR1 and CCR3 and induction of chemotaxis to CC chemokines by IFN-γ in human neutrophils. J Immunol, 162, 474–479.

